# The neural time course of size constancy in natural scenes

**DOI:** 10.1101/2024.09.26.615104

**Authors:** Lu-Chun Yeh, Surya Gayet, Daniel Kaiser, Marius V. Peelen

## Abstract

Accurate real-world size perception relies on size constancy, a mechanism that integrates an object’s retinal size with distance information. The neural time course of extracting pictorial distance cues from scenes and integrating them with retinal size information - a process referred to as scene-based size constancy - remains unknown. In two experiments, participants viewed objects with either large or small retinal sizes, presented at near or far distances in outdoor scene photographs, while performing an unrelated one-back task. We applied multivariate pattern analysis (MVPA) to time-resolved EEG data to decode the retinal size of large versus small objects, depending on their distance (near versus far) in the scenes. The objects were either perceptually similar in size (large-near versus small-far) or perceptually dissimilar in size (large-far versus small-near), reflecting size constancy. We found that the retinal size of objects could be decoded from 80 ms after scene onset onwards. Distance information modulated size decoding at least 120 ms later: from 200 ms after scene onset when objects were fixated, and from 280 ms when objects were viewed in the periphery. These findings reveal the neural time course of size constancy based on pictorial distance cues in natural scenes.

## Introduction

Scene context provides information about the distance and thereby the real-world size of objects, facilitating object recognition, and constraining the ways we can interact with objects. While we can retrieve distance information from stereopsis, which is especially useful for nearby objects, in natural vision we strongly rely on pictorial distance cues such as the relative size of objects, height in the visual field, occlusion, image blur, and linear perspective (Cutting & Vishton, 1995). These pictorial distance cues are combined with the retinal size of objects to infer the real-world size of these objects, a process often referred to as the size constancy mechanism (Holway and Boring, 1941; Gruber, 1954; for a review, Sperandio and Chouinard, 2015). The importance of linear perspective in size constancy is demonstrated by the famous Ponzo illusion: two objects with identical retinal sizes are perceived as having different sizes when presented on the upper and lower sides of a pair of converging lines (Ponzo, 1911), presumably because the converging lines are interpreted as parallel lines (e.g., train tracks) extending away from the observer.

Several studies have provided neural evidence that pictorial distance cues modulate visual cortex responses to the same stimulus placed near or far in a scene. For example, human fMRI studies showed that stimulus representations in the primary visual cortex (V1) are affected by pictorial distance information (Murray et al., 2006; Fang et al., 2008; Schwarzkopf et al., 2011; He et al., 2015). In these studies, a variant of the Ponzo illusion was used to vary perceived size while keeping the retinal size constant. Results showed that objects that were supposedly perceived as larger, due to their position in the scene (far away), activated a larger part of V1. In line with these findings, an electrophysiological study in non-human primates showed that the firing profile of V1 neurons reflected the rescaled size based on contextual distance information (Ni et al., 2014). However, the cortical location of size constancy does not speak to the time course of size constancy; modulations in V1 could occur both early and late in time, for instance due to feedback projections from object- or scene-selective regions (Zeng et al., 2020).

Previous studies investigating the neural time course of size constancy displayed the scene context (with pictorial distance cues) *before* the objects appeared at the near of far location of the scene. In these studies, a differential neural responses to objects placed far versus near in a scene occurred during the initial evoked response in monkey V1 (<100 ms; Ni et al., 2014), and shortly after the initial evoked response (at around 150 ms) in human EEG (Chen et al., 2019). As the distance cues were already presented before stimulus onset in these studies, however, expectations of object size (i.e., a rescaling factor) may have been established before stimulus onset. It therefore remains unknown how retinal size and pictorial distance cues are combined to generate a representation of object size when perceiving objects and scenes simultaneously, as is often the case during day-to-day perception. Thus, in the present study, we investigated the time course of size constancy from the moment of scene (and object) onset.

We hypothesized that size constancy would emerge relatively late during object processing, as it would require extracting distance cues from the scene first. For example, previous MEG studies showed that facilitatory interactions between scene and object processing in recognition tasks (e.g., a road disambiguating the representation of a car) occur relatively late, around 300 ms after stimulus onset (Brandman & Peelen, 2017; 2023; Leticevscaia et al., 2024). Accordingly, scene-based size constancy may similarly arise from around 300 ms after scene onset. Alternatively, size constancy may start earlier (perhaps as early as the initial evoked size response; Ni et al., 2014), considering that distance cues (e.g., linear perspective) are consistent across scenes, and that size-distance scaling applies generally across objects, unlike the more specific semantic relations between scene and object categories. In line with such an early size-consistency mechanism, it was shown that briefly presented objects in scenes attract attention when their perceived size matches a memory template, compared with perceptually mismatching objects subtending the same retinal size (Gayet & Peelen, 2019).

Here, we set out to test the neural time course of scene-based size constancy in humans, testing when size representations transfer from proximal (i.e., retinal image size) to distal (i.e., perceived real-world size). In two experiments, we employed a multivariate pattern analysis (MVPA) approach to decode size information from time-resolved EEG response patterns. Participants viewed objects subtending large or small retinal sizes, presented in the near or far plane of natural scenes, while performing an unrelated one-back task. We decoded the retinal size of the objects (large versus small) over time, depending on their viewing distance within the scene (near versus far). Specifically, we compared the decoding time course of large versus small objects that were either perceptually relatively similar in size (large-near versus small-far) or perceptually relatively different in size (large-far versus small-near), due to size-distance rescaling. This analysis approach allowed us to quantify how representations of object size get transformed across the visual processing cascade, from representations of retinal size to representations of perceived real-world size.

## Materials and Methods

### Participants

The target sample size (N = 36) for each experiment was determined by reaching 80% power for detecting a hypothetic medium effect of d =0.5 (N = 34) and counterbalancing four sets of stimuli between participants. Thirty-six healthy volunteers (26 females, mean age = 23.14, SD = 3.036 years) participated in Experiment 1. For Experiment 2, we could only recruit 31 volunteers (23 females, mean age = 22.68, SD = 4.21 years) within the testing period allocated to the project. All participants had normal or corrected-to-normal vision. The participants gave informed written consent and received a gift card of 10 euro per hour for their time. The study was approved by the Ethics Committee of the Faculty of Social Sciences, Radboud University Nijmegen.

### Materials and Resources

All materials and data are publicly accessible via the following online repository: https://osf.io/pxs7w/

### Design and Stimuli

We manipulated the retinal size of objects (large and small) and their distance within scenes (near and far) to create 4 types of stimuli (Figure 1A). The full stimulus set (for the main experiments) comprised twelve object categories (six animate and six inanimate object categories), and 64 outdoor scenes (48 scenes for the formal experiment and 16 for the practice trials). There were 12 exemplars in each object category; four of those were combined with one scene each, and the remaining eight were presented in isolation on a gray background (i.e., the isolated object condition; see below). Each object-scene pair generated 4 types of stimuli (large-near, large-far, small-near, and small-far), but each participant only saw each scene-object pair in one of the types, yielding four stimulus sets that were counterbalanced across the participants. The objects presented within scenes were degraded, impairing object recognition, to mitigate the influence of participants’ prior knowledge of the objects’ real-world sizes. This design also allowed us to examine whether size-distance congruency facilitates object recognition but these results are not reported here. All scene-object stimuli were reduced to 355 (horizontal) x 266 (vertical) pixels, subtending about 10° * 7.5° of visual angle, and were presented on a gray background. In Experiment 1, scene-object stimuli were presented at the center of the screen. In Experiment 2, the objects within the scenes were presented at fixation, inevitably causing the scenes to vary in position across trials. Because the variable position of the scenes induced large trial-by-trial variation in visual input, low-level visual stimulation across the presentation area was equated by placing the stimuli on top of a larger phase-scrambled background, 800 (horizontal) x 600 (vertical) pixels (Figure 1B).

**Figure 1.**
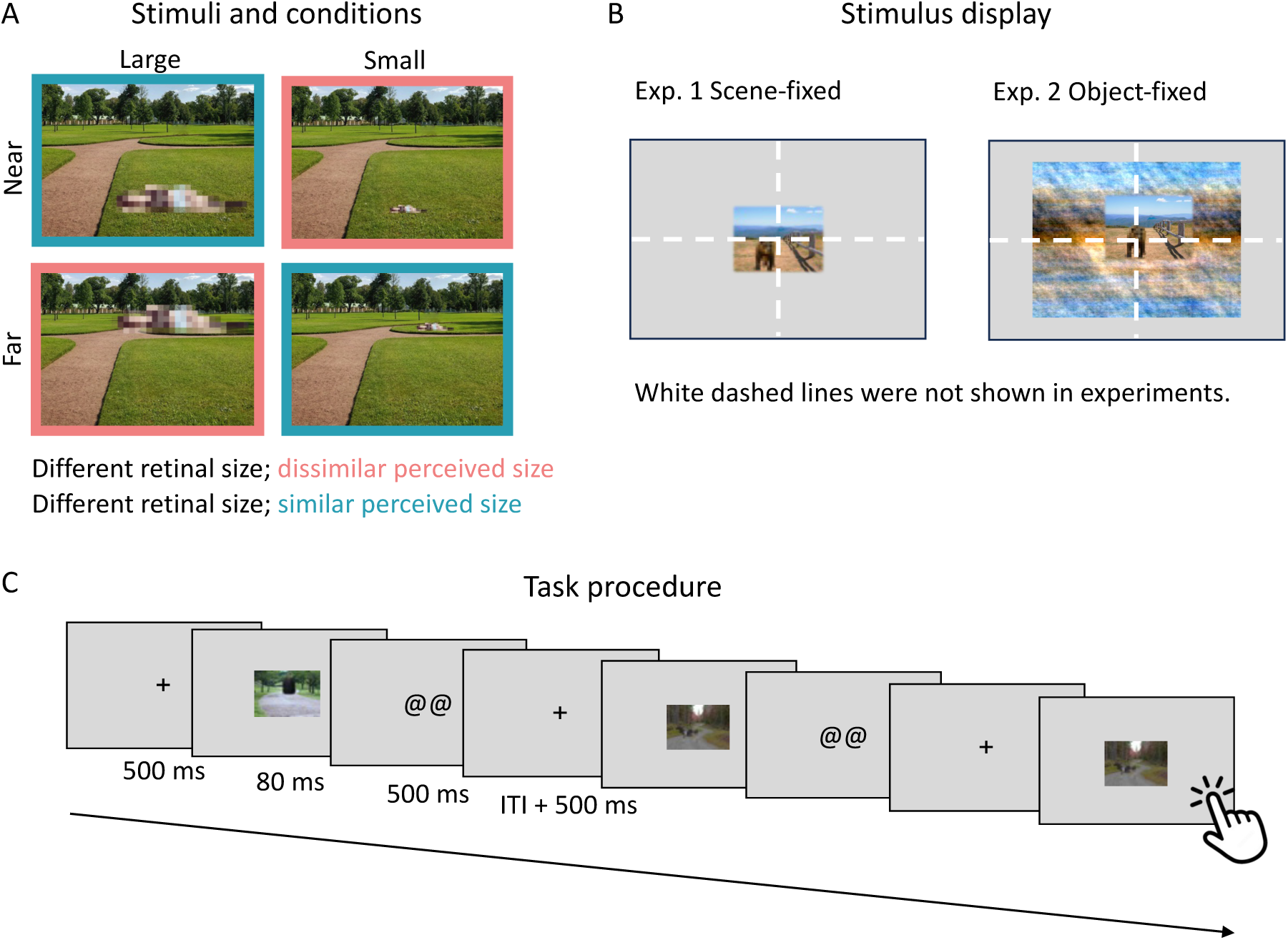
Stimuli and task. (A) Four types of stimuli creating the two conditions of interest (dissimilar perceived size; here, outlined in red, similar perceived size; outlined in blue). (B) Stimulus displays of Experiments 1 and 2. In Experiment 1, the scenes were always centered at fixation, so that the location of objects varied across trials. In Experiment 2, the objects were always centered at fixation, so that the location of the background scenes varied across trials. Scrambled backgrounds were used to balance the visual input across the visual field. (C) Task procedure of the one-back task. Participants were instructed to fixate the central cross, and press the space bar when the same stimulus was presented twice in succession (ITI: intertrial interval).

### Experiment Procedure

EEG data were recorded while participants performed a one-back repetition task. Participants sat in an illuminated room at a distance of 57 cm from the monitor. First, participants received instructions about task requirements and practiced two blocks (one with degraded objects within natural scenes and one with intact isolated objects presented on the gray background). Each practice block contained 54 trials, including six trials in which an image was presented twice in succession, and a response was expected (i.e., one-back task). The stimuli used in the practice block were different from the stimuli used in the formal experiment. The formal experiment consisted of 12 blocks of 108 trials, including 12 trials with stimulus repetitions (hereafter: repetition trials). Eight blocks comprised objects within natural scenes (as described above), and four blocks comprised objects from the same categories (different exemplars, but with similar viewing angles and postures) presented at fixation on top of a gray background. The order of blocks was randomized and varied between participants. Each trial started with a 500-ms fixation cross presented at the center of the screen. Then, a stimulus was presented for 80 ms and followed by another fixation for 800 ms. After that, a 500-ms slide with “@@” was presented to remind participants that they can blink during this period. The intertrial intervals varied randomly from 500 to 900 ms with a mean of 700 ms. The task procedure is illustrated in Figure 1C.

Participants were instructed to fixate on a cross at the center of the screen throughout the practice and experimental blocks, and to respond as quickly and accurately as they could whenever they detected a stimulus repetition. Participants were also instructed to minimize eye blinking and movement during stimulus presentation and to only blink during the specified time window (the 500-ms slide with “@@”). Participants could take a rest between blocks, and they could self-initiate the next block. The full experimental time for each participant was around 60 minutes.

### EEG Data Acquisition

Electrophysiological data were recorded using a customized 64-channel active electrode actiCAP system with 500 Hz sampling rate. The electrodes included seven sites in the central line (Fz, FCz, Cz, CPz, Pz, POz, and Oz) and 28 sites over the left and right hemispheres (FP1/FP2, AF3/AF4, AF7/AF8, F1/F2, F3/F4, F5/F6, F7/F8, FC1/FC2, FC3/FC4, FT5/FT6, FT7/FT8, FT9/FT10, C1/C2, C3/C4, C5/C6, T7/T8, CP1/CP2, CP3/CP4, CP5/CP6, TP7/TP8, P1/P2, P3/P4, P5/P6, P7/P8, PO3/PO4, PO7/PO8, PO9/PO10, and O1/O2). AFz served as the ground electrode, and TP9/TP10 placed on the left/right mastoid as reference electrodes. Vertical and horizontal eye movements were also recorded using EOG electrodes. Impedance of all the electrodes was kept below 20 kΩ. Trial-type codes were sent to the computer used to record EEG data via a parallel port from the computer used to present task stimuli.

### EEG Data Analyses

#### Preprocessing

EEG data were preprocessed offline using the Fieldtrip toolbox (Oostenveld et al., 2011) in MATLAB (MathWorks). To remove artifacts caused by eye movements and blinks, an independent component analysis was performed first, and a visual inspection was used to identify and remove the eye movement and blink components from the data of each participant. Then, EEG data were re-referenced using the average of all electrodes except EOG, FP1, and FP2, and epoched from -100 to 500 ms relative to stimulus onset. Epochs were baseline-corrected using the prestimulus period (from -100 to 0 ms). Epochs of repetition trials were excluded from further analyses.

#### Decoding object size in scenes

Multivariate classification analyses were conducted using the Amsterdam Decoding and Modeling toolbox (ADAM; Fahrenfort et al., 2018). Linear discriminant analysis was used to decode objects’ retinal size (large versus small). To focus on the brain regions involved in visual processing, only data from posterior electrodes were used in the analyses (CPz, Pz, POz, Oz, T7/T8, CP1/CP2, CP3/CP4, CP5/CP6, TP7/TP8, P1/P2, P3/P4, P5/P6, P7/P8, PO3/PO4, PO7/PO8, and O1/O2). The data were resampled to 100Hz, so classification was performed for every 10 ms. The Area Under the Curve (AUC) was used as the measure of decoding sensitivity; higher AUC reflects better discrimination between classes.

We created two experimental conditions, from the four possible stimulus combinations (Figure 1A): (1) In the similar perceived size condition, classifiers were trained and tested on data evoked by stimuli with large-near vs. small-far objects, which have different retinal sizes and relatively similar perceived sizes when distance information is taken into account. (2) In the dissimilar perceived size condition, classifiers were trained and tested on data evoked by stimuli with large-far vs. small-near objects, which have equally different retinal sizes, but the perceived sizes are drastically different when distance information is taken into account.

Size classification for both conditions was done using a 10-fold cross-validation procedure. The trials of the relevant conditions were divided into 10 equally-sized folds. Then, a leave-one-out procedure was used; 9 folds were used as training data, and the left-out fold was used as testing data. This procedure was repeated 10 times, iterating through all 10 folds, and classifier performance was then averaged over these 10 folds (see Figure 2A for the cross-validation analysis schematic). The analysis was conducted using a cross-temporal approach, yielding all possible combinations of 10ms timepoint pairs for training and testing the classifiers. After classification analyses, we used a cluster-based nonparametric permutation one-sample *t*-test (Maris & Oostenveld, 2007) to determine the significant time window of above-chance classification while correcting for multiple comparisons. First, a *t*-test was run for each time point; then, if there were continuous time points whose *t-*values were larger than the threshold, they were clustered together. The sum of the *t*-values within clusters was computed, and their probability was compared to that of a null distribution created from 1000 Monte Carlo random partitions, and cluster correction for type I error (*p* <.05, one-tailed). Finally, we tested the difference in decoding performance between conditions (more versus less similar perceived size difference) using a permutation-dependent *t*-test (two-tailed) to correct for multiple comparisons.

**Figure 2.**
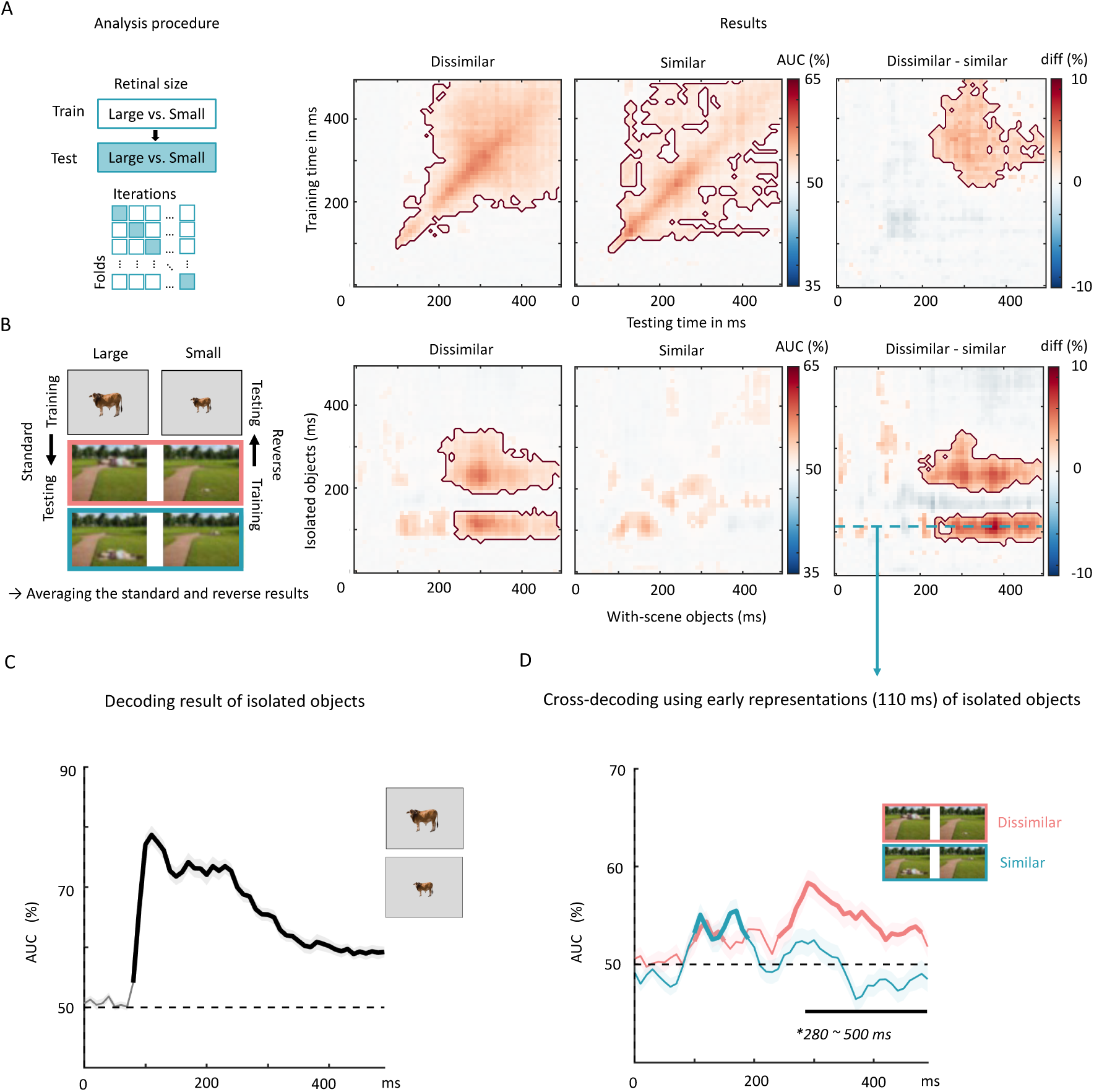
Results of the scene-fixed experiment (Exp. 1). (A) Procedure and results of the cross-validation decoding analysis. We trained and tested classifiers to decode the retinal size of degraded objects presented within scenes, separately for the similar and dissimilar perceived size conditions. The trials for each condition were divided into 10 equally-sized folds. The results matrices represent the AUC across time from 0 to 500 ms after stimulus onset. The two left-most matrices show the AUC of the two conditions separately. The black outline demarcates significant clusters of decoding performance for individual conditions (*p* <.05, one-tailed). The right matrix shows the difference between dissimilar and similar conditions. The black outline demarcates significant clusters of decoding performance differences between the two conditions (p <.05, two-tailed). (B) Cross-decoding analysis procedure and results. We trained classifiers to distinguish between large and small (isolated, non-degraded, centrally presented) objects on a gray background, to decode the size of (degraded, variably positioned) objects within scenes. We also used the same data, but reversing the train-test direction (training on objects within scenes, and testing on isolated objects), and averaged the results across these two train-test directions. Following the visualization principles of panel 2A, the two left-most matrices show the AUC of the two individual conditions, whereas the right matrix shows the difference in decoding performance between conditions. (C) Results of the analysis decoding the retinal size (large vs small) of isolated objects. (D) Results of a cross-decoding analysis using early representations (110 ms, see the blue dashed lines in panel B) of isolated objects to decode the size of objects in scenes (throughout the entire epoch). Significant clusters for each condition are highlighted in bold (*p* <.05, one-tailed). The black-bold line indicates significant differences between the two conditions (*p* <.05, two-tailed).

#### Cross-decoding object sizes in scenes and in isolation

In addition to the decoding analysis described above, we ran a cross-decoding analysis to isolate the neural responses to object size and to exclude contribution of other features that may differ between the similar and dissimilar perceived size conditions. To achieve this, classifiers were trained to distinguish between large and small objects presented in isolation on a gray background (i.e., without any distance information), and tested to distinguish between large and small objects within scenes. This analysis was also conducted in the opposite train-test direction (training on objects in scenes, and testing on isolated objects), and results were averaged across these directions. Figures 2A and 3A illustrate this cross-decoding procedure. We also ran the cross-decoding analysis with a cross-temporal approach. This approach reveals the dynamic change of object-size representations, in the presence and absence of scene context. After performing these classification analyses, we used a cluster-based nonparametric permutation one-sample *t*-test (Maris & Oostenveld, 2007) to determine the significant time window of above-chance classification, and tested the difference in decoding performance between the more and less similar perceived size conditions using the permutation dependent *t*-test (two-tailed).

**Figure 3.**
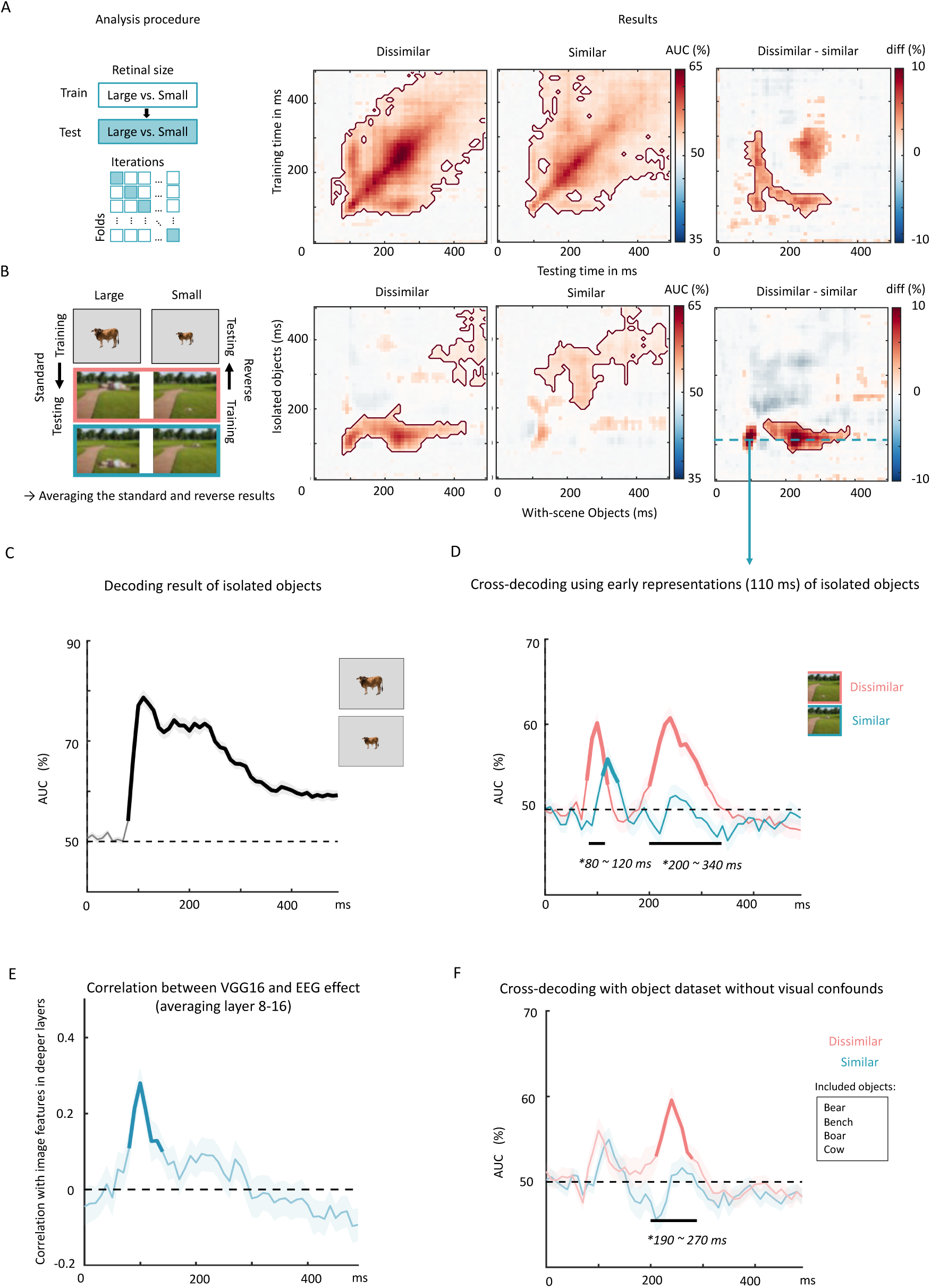
Results of the object-fixed experiment (Exp. 2). (A) Procedure and results of the cross-validation decoding analysis, following the procedures of Experiment 1. The results matrices represent the AUC across time from 0 to 500 ms after stimulus onset. The left and middle matrices show the AUC of the two conditions separately. Significant clusters for each condition are highlighted in red-bold line (*p* <.05, one-tailed). The right matrix shows the difference between dissimilar and similar perceived size conditions. The red-bold lines show the significant clusters (*p* <.05, two-tailed). (B) The cross-decoding analysis procedure and results. We trained classifiers to distinguish between large and small (isolated, non-degraded, centrally presented) objects on a gray background, to decode the size of (degraded, variably positioned) objects within scenes. We also used the same data, but reversing the train-test direction (training on objects within scenes, and testing on isolated objects), and averaged the results across these two train-test directions. Results were averaged across these directions. The left and middle matrices show the AUC for the two similarity conditions. Significant clusters for each condition are highlighted in red-bold line (*p* <.05, one-tailed). The right matrix shows the difference between dissimilar and similar perceived size conditions. The red-bold lines show the significant clusters (*p* <.05, two-tailed). (C) Results of the cross-validation analysis decoding the retinal size of isolated objects on a gray background. (D) Results of the cross-decoding analysis using early representations (110 ms, see the blue dashed lines in panel B) of isolated objects. Significant clusters for each condition are highlighted in bold (*p* <.05, one-tailed). The black-bold line indicates significant differences between the two similarity conditions (*p* <.05, two-tailed). (E) Results of the correlation analysis correlating dissimilar-similar differences in VGG16 and EEG. We first computed the Pearson correlation between each subject’s EEG data and the VGG16 data for the images that they saw during the experiment. Then, we tested the correlation coefficients from 31 participants against chance using a permutation *t*-test (*p* <.05, two-tailed). Results showed significant cluster of correlation between EEG data and VGG16 image analysis data (bold line) from 80-120 ms. (F) The results of cross-decoding analysis using early representations (110 ms) of isolated objects, using only those four object categories yielding no visual confounds according to the VGG16 image analysis (see Supplementary Figure 2A for details). Significant clusters for each condition are highlighted in bold (*p* <.05, one-tailed). The black-bold line indicates significant differences between the two conditions (*p* <.05, two-tailed).

## Results

### Experiment 1: Scenes presented at fixation

In Experiment 1, scenes were centered at fixation, such that the objects would appear either above (far) or below (near) fixation. Behavioral performance in the one-back repetition task was near ceiling (96.81% ± 1.76%), confirming that participants followed our instructions and paid attention to the stimuli.

To test how distance cues modulated object size representations in scenes, we trained and tested classifiers to decode retinal object size (large vs small) using degraded objects within scenes, separately for both conditions of interest (similar perceived size, dissimilar perceived size). This was done using a temporal generalization approach. The results are illustrated in Figure 2A. Retinal size could be decoded from EEG patterns from around 90 ms after stimulus onset onwards in both conditions. To investigate the timing of size constancy, we tested the difference in decoding performance between the two conditions. On the diagonal, decoding performance in the similar perceived size condition was significantly lower than in the dissimilar perceived size condition between ∼280 to 390 ms after stimulus onset. The significant difference between similar and dissimilar perceived size conditions starting at 280 ms indicates that pictorial distance cues modulated representations of retinal object size. Moreover, this size constancy mechanism emerged only after about 280 ms, which is substantially later than the emergence of size representations per se.

In the decoding analyses described above, it is unclear which features contributed to the decoding. In principle, any feature that differed between the similar and dissimilar perceived size conditions could drive the observed decoding effects. For example, the locations of the large and small objects differed in the two conditions (large object above vs below fixation). Therefore, to isolate the influence of the scenes on size processing, we turned to cross-decoding analyses, using an independent data-set for training the classifier: the runs in which participants viewed isolated, centrally presented, large and small objects on a gray background.

In the cross-decoding analysis, classifiers were trained to decode the retinal size of isolated objects (at fixation) and were then tested to decode the retinal size of objects in scenes (at varying positions). The reverse cross-decoding direction analysis was also conducted; the reported results are the average of the two directions (Man et al., 2012; Oosterhof et al., 2010). The key aim was again to test for decoding differences between similar and dissimilar perceived size conditions. Cross-decoding results are illustrated in Figure 2B. Contrasting decoding performance between the two conditions revealed two significant clusters (Figure 2B, right panel). The first of these clusters showed that early size representations of isolated objects (90-150 ms after stimulus onset) generalized better to dissimilar than to similar perceived size conditions from 240 ms after stimulus onset. The second cluster centered around relatively late representations of isolated objects (220-300 ms after stimulus onset). These representations also generalized better to dissimilar than to similar perceived size conditions but this difference reached significance slightly earlier in time (from around 200 ms; Figure 2B, right panel).

Of the two clusters, the cluster centered around the early-stage representation of isolated object size is of particular interest for addressing our research question, as such early size representations are most likely to reflect visually-driven (retinal) size representations. By contrast, later-stage representations of isolated object size will additionally include high-level object information, such as the implied distance of the object given its retinal size, complicating the interpretation of the effects we observed in the cross-decoding analyses. Therefore, to address the question of how representations of retinal size were modulated by pictorial distance cues, we ran an additional targeted analysis focusing on the early representations of isolated object size. For this analysis, we first ran an analysis to decode retinal size within the isolated object runs to establish the first peak latency at 110 ms (Figure 2C). Then, we fixed the training/testing time of the isolated object data on this time point and tested for generalization to the main experiment data. The reverse cross-decoding direction analysis was also applied and averaged with the analysis results obtained with the standard decoding direction. This analysis revealed that the retinal size of objects in scenes could be decoded from 100 to 200 ms in the similar condition and from 100 to 160 ms, 170 to 210 ms, and 240 to 490 ms in the dissimilar condition. Importantly, the difference between the similar and dissimilar conditions was again significant from 280 ms after stimulus onset (Figure 2D).

#### Interim summary

Experiment 1 showed that retinal size representations could be decoded from around 90 ms after stimulus onset but were only modulated by pictorial distance cues (reflecting size constancy) from around 280 ms after stimulus onset. These findings are consistent with previous studies that found that interactions between scene and object processing occur around 280-320 ms after stimulus onset (Brandman & Peelen, 2017, 2023; Leticevscaia et al., 2024; Wischnewski & Peelen, 2021).

### Experiment 2: Objects presented at fixation

Experiment 1 showed that size constancy started relatively late. However, in this experiment, objects appeared away from fixation and at varying locations across trials. Combined with the fact that objects were not directly relevant for the task, it is possible that objects were poorly attended and/or poorly visible, delaying (or watering down) modulatory influences. Therefore, in Experiment 2, the scenes were presented at varying locations within the presentation area, in such a way that the objects always appeared at fixation. In this case, both far and near objects appeared at the exact same retinal location, and were always foveated (and presumably attended) during the 80 ms presentation duration. This should drive EEG responses to the objects more strongly, more systematically, and more quickly. Furthermore, this design avoids possible confounds following from the difference in visual field stimulation by far and near objects (i.e., object above vs below fixation). Finally, placing the objects at fixation increases their representational overlap with the objects in the isolated object condition, presumably improving cross-decoding performance. Below, we report the same analyses as reported for Experiment 1. Behavioral performance in the one-back repetition task was again near-ceiling (97.03% ± 1.85%.), confirming that participants followed our instructions and paid attention to the stimuli.

We trained and tested classifiers on EEG data to decode the retinal size of objects within scenes, separately for both perceived size conditions (similar perceived size, dissimilar perceived size). The results are illustrated in Figure 3A. In both conditions, retinal object size could be decoded from EEG patterns from around 80 ms after stimulus onset (p <.001). To reveal the time course of size constancy, we again tested the difference in decoding performance between the two conditions. Unlike Experiment 1, we found a significant cluster showing that early size representations (around 80-150 ms after stimulus onset) were decoded better for the dissimilar than the similar condition (p =.024), suggesting a very fast size constancy mechanism (i.e., emerging simultaneously with size representations per se). We more closely investigate this early effect in subsequent analyses below.

In the cross-decoding analysis, classifiers were trained to distinguish between large and small isolated objects, and were tested on their ability to distinguish between large and small objects within scenes. This analysis was also conducted in the reverse train-test direction, and the reported results are the average of the two decoding directions. Cross-decoding results are illustrated in Figure 3B. Contrasting decoding performance between the dissimilar and similar perceived size conditions revealed a significant cluster in which the early representation of isolated object size (80-170 ms after stimulus onset) better generalized to the dissimilar than the similar perceived size condition, starting from around 140 ms after stimulus onset (Figure 3B, right panel), and thus substantially earlier than in Experiment 1.

Finally, as in Experiment 1, we ran an additional targeted analysis focusing on the early representations of retinal object size, based on activity evoked by viewing large versus small isolated objects. As in Experiment 1, peak decoding for isolated objects was observed at a latency of 110 ms (Figure 3C). Then, using this time point to obtain early representations of (isolated) object size, we tested for generalization to the main experiment data (objects in scenes). This analysis revealed that retinal object size could be decoded from 110 to 150 ms in the similar condition and from 80 to 130 ms and 200 to 320 ms in the dissimilar condition. The difference between the similar and dissimilar conditions was significant from 200 to 340 ms (Figure 3D), and thus slightly earlier than that of Experiment 1A. Additionally, we found a significant early cluster from 80 to 120 ms, suggesting a very fast size constancy mechanism, emerging as early as initial retinal size representations.

#### Image-similarity analysis

Considering the surprisingly early onset of size-constancy in Experiment 2, we wondered whether this result could be explained by uncontrolled differences in local image features between the dissimilar and similar perceived size conditions. While the large and small objects were identical in the two conditions, the local background differed. That is, in the dissimilar perceived size condition, the small objects were presented on near backgrounds and the large objects on far backgrounds, and the opposite was true for the similar perceived size condition. It is possible, in principle, that image features of the local background (e.g., spatial frequency, luminance contrast) contributed to size decoding when the object (and its local background) was foveated, as was the case in Experiment 2. This cannot be ruled out even in the cross-decoding analysis if the local backgrounds happen to share features with large or small isolated objects (e.g., if near backgrounds share features with small isolated objects, this would benefit decoding in the dissimilar perceived size condition where small objects were presented on near backgrounds). To investigate this possibility, we ran an image similarity analysis using a deep neural network (VGG16; see Supplementary Material for a full report of this analysis). Results showed that image features of a cropped segment centered on the object (visual angle = 3 degrees) indeed provided information about the distance-inferred size of the objects. Specifically, image features that differentiated large versus small isolated objects more closely resembled image features differentiating large versus small objects in the dissimilar than the similar perceived size comparison (Supplementary Figure 1). Conducting this analysis on individual scenes showed that this may have affected about half the scenes (Supplementary Figure 1). To test whether these image-similarity results could explain the (early) EEG decoding results, we correlated the dissimilar-similar differences in VGG16 with the corresponding differences in EEG across individual scenes. Results showed a positive correlation for the early cluster (80-120 ms; *r*=0.52; *p* <.05, one-tailed) but not the late cluster (200-340 ms; *r*=0.046; *p*=. 44, one-tailed). This result was confirmed in a correlation analysis across the whole EEG decoding time course, showing that the VGG16-EEG correlation was specific to the early decoding difference (Figure 3E). In a final analysis, we reran the EEG decoding analysis after excluding those images that showed image-level differences between dissimilar and similar perceived size conditions (as established using VGG16). This analysis showed that the early cluster disappeared when controlling for image-level differences (Figure 3F). Importantly, however, the later cluster remained strong and significant, from 190-270 ms after scene onset.

#### Interim summary

In the object-fixed experiment (Experiment 2), we again found evidence for size constancy effects across a set of complementary analyses. Results showed that retinal size representations could be decoded from around 80 ms after stimulus onset and that these were modulated by pictorial distance cues (reflecting size constancy) from around 200 ms after stimulus onset. An even earlier modulation of size decoding was shown to be related to differences in image characteristics between stimulus conditions, and disappeared when controlling for these differences.

## Discussion

In this study, we investigated the time-course of scene-based size constancy: when size representations shift from reflecting the retinal size to the perceived size of an object, based on pictorial distance cues. To address this, we used multivariate pattern analysis of EEG data, where we decoded the size of objects within scenes. By contrasting conditions that were perceived as having relatively dissimilar versus similar size, we could delineate when representations shift towards the coding of perceived size.

Across two experiments, results showed a stable and strong representation of perceived size that emerged from 200 ms after stimulus onset. This was substantially later than the emergence of size representations per se, which we observed from 80-90ms onwards and peaked at 110ms after stimulus onset. The time course of size constancy revealed here is also later than that of previous studies, which reported size constancy at or before 150 ms relative to object onset (Ni et al., 2014; Chen et al., 2019; Zeng et al., 2020). The main difference between these previous studies and ours is that, in these previous studies, the distance cues were already processed before the object was presented. For example, in Ni et al. (2014), the background (with linear perspective) was presented throughout a block, and size constancy was time-locked to the onset of the object that was presented on top of the background. Therefore, their time course does not include the processing of distance cues from the scene, as in our study. Similarly, in Chen et al. (2019), the monitor was moved close or far from the observer before stimulus onset. Here as well, then, the time course of size constancy did not include processing of distance cues. Indeed, recent work suggests that observers may actively account for viewing distance by preemptively rescaling object representations (Gayet, & Peelen, 2022; Gayet et al., 2024). However, in day-to-day vision, objects typically do not appear within pre-viewed scenes; whenever a region of the visual field is fixated after a saccade, we have to simultaneously process objects and their surrounding scene.

The scene-based size constancy mechanism investigated here involved extracting pictorial distance cues from scenes and modulating representations of object size based on these cues. Accordingly, we propose that scene-based size constancy starts by representing scene layout, in scene-selective cortex (Kravitz et al., 2011; Park et al., 2011; Henriksson et al., 2019), which then modulates representations in object-selective cortex. Subsequently, object-selective cortex may feed back to early visual cortex, as revealed by a TMS study investigating size constancy (Zeng et al., 2020), ultimately leading to changes in perceived size. In their study, TMS to object selective cortex selectively reduced size constancy effects during earlier time periods than TMS to early visual cortex, suggesting that the effects in early visual cortex were driven by feedback from object-selective areas. Although there is no direct evidence yet for the involvement of scene-selective regions in the size constancy mechanism, our findings – together with those of Zeng and colleagues (2020) – demonstrate the role of feedback in the distance-dependent rescaling of early visual representations. This stands in contrast with the view that size constancy emerges jointly with early size representations in early visual cortex (Ni et al., 2014).

The exact timing at which pictorial distance cues started affecting visual size representations differed between our two experiments (even after controlling for low-level visual differences). These experiments differed in the presentation location of the scenes, with presentation either centered on the scene, with the object appearing in unpredictable and peripheral locations (Experiment 1), or with presentation centered on the object, such that the object was foveated (Experiment 2). Size constancy occurred earlier in the object-fixed experiment (200 ms) than in the scene-fixed experiment (280 ms). This difference may reflect the additional time needed for object localization when object location is unpredictable. It may also reflect the weaker and more variable responses evoked by peripherally presented objects. The time course of the scene-fixed experiment is in line with previous MEG studies investigating the influence of scene context on object category representations (Brandman & Peelen, 2017; Leticevscaia et al., 2024). In these studies, objects similarly appeared at varying and unpredictable locations in scenes. Similar to the current findings, scene context facilitated object category decoding from 280 ms after scene onset. This raises the possibility that these contextual effects on representations of object size (measured here) and on representations of object category reflect a common mechanism, relying on the same interaction between scene and object processing regions. Alternatively, the effects of scene context on object category representations may be (partly) attributed to the effects of pictorial distance cues, which disambiguate the real-world size of objects. In line with this idea, it was shown that objects are better recognized and detected when their retinal size matches rather than mismatches their viewing distance in the scene (Biederman, et al., 1982; Eckstein et al., 2017; Wolfe, 2017; Gayet et al., 2022).

*Conclusion.* In summary, our results reveal the time course of object size perception in natural scenes. An object’s retinal size was initially represented around 80-90 ms after stimulus onset. These retinal size representations were modulated by the object’s location in a scene, starting approximately 200 ms after stimulus onset when the object was fixated. This time likely reflects the need to first extract distance cues from the scene and then integrate these with retinal size. Overall, our results provide insight into how the visual system transforms retinal size representations into the behaviorally adaptive perception of real-world object size.

## Acknowledgements

The project received funding from the European Research Council under the European Union’s Horizon 2020 research and innovation program (ERC; grant agreement no. 725970, granted to M.V.P.).

