## Supplementary Material for "The neural time course of size constancy in natural scenes"

### Supplementary Materials: image similarity analysis

*Supplementary materials accompanying the study entitled "The neural time course of size constancy in natural scenes" by Lu-Chun Yeh, Surya Gayet, Daniel Kaiser, and Marius V. Peelen.*

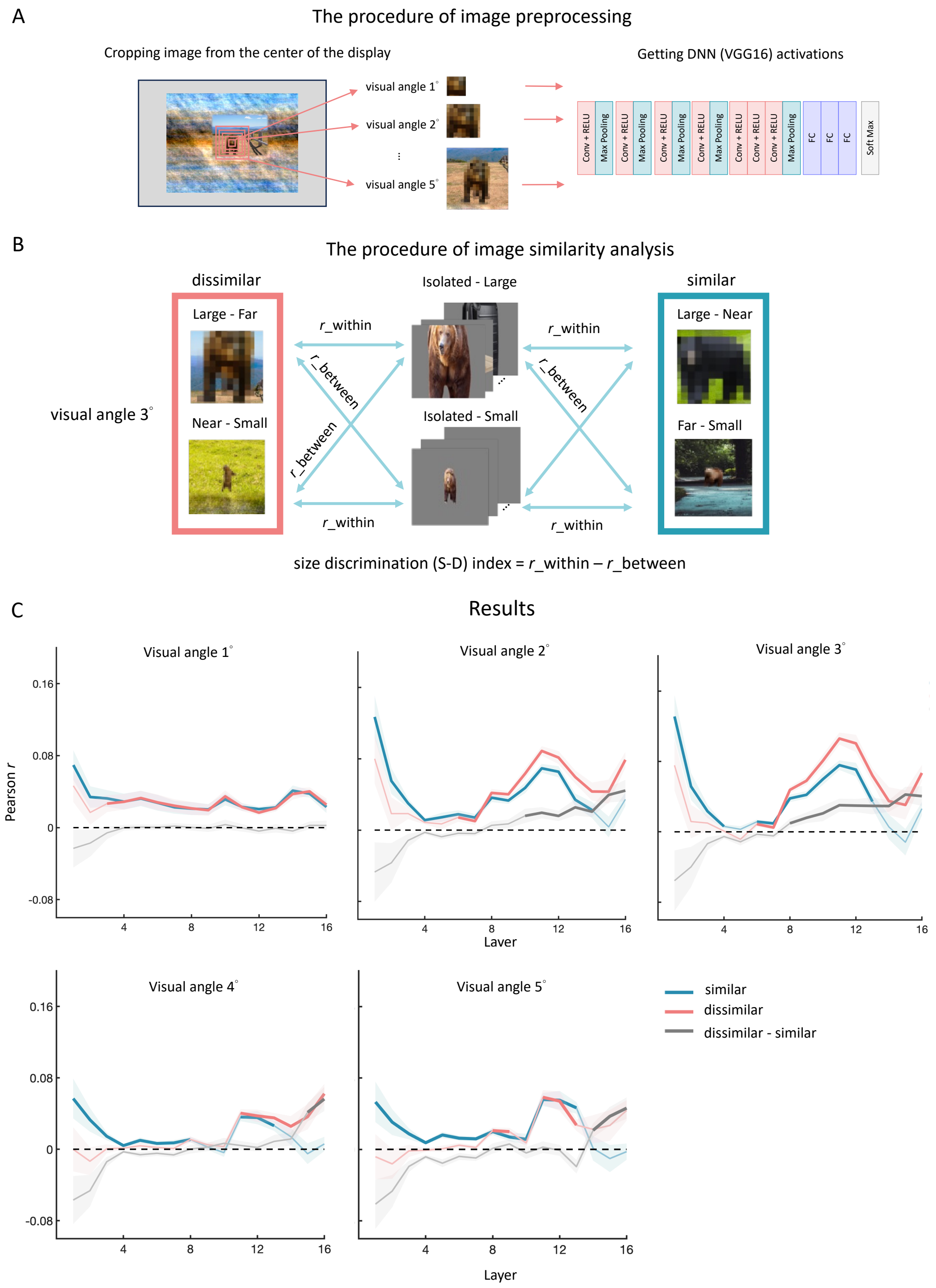

**Supplementary Figure S1.** Results of image similarity analysis for the object-fixed experiment (Experiment 2). (A) The image processing procedure. Images were cropped from the center of the display using a rectangular aperture (sides of 1, 2, 3, 4, and 5 degrees of visual angle). The cropped images were then processed through a deep neural network model, VGG16, to obtain layer-wise activations for each image in the stimulus set used in the EEG experiments. (B) Image similarity analysis procedure. Correlations were calculated between each cropped image from similar and dissimilar perceived size conditions and the isolated object images (96 images per size) across all layers. The average correlation coefficients were determined across images, generating one "within-size" correlation (i.e., images of the same size, with and without a scene) and one "between-size" correlation (i.e., images of different size, with and without a scene) per layer. The size discrimination index was computed by subtracting the between-size correlation from the within-size correlation. (C) Results of the image similarity analysis. We first tested the size discrimination index against zero using a permutation *t*-test ($p < .05$, one-tailed). Then, we tested the difference between similar and dissimilar perceived size conditions using permutation paired *t*-tests ($p < .05$, two-tailed). Significant clusters for each condition and their difference are highlighted in bold. Generally, the size discrimination index was larger in the dissimilar perceived size condition than the similar perceived size condition in the later layers of VGG16 (using apertures of around 3 degrees), which is reminiscent of the EEG cross-decoding results reported in the main manuscript.

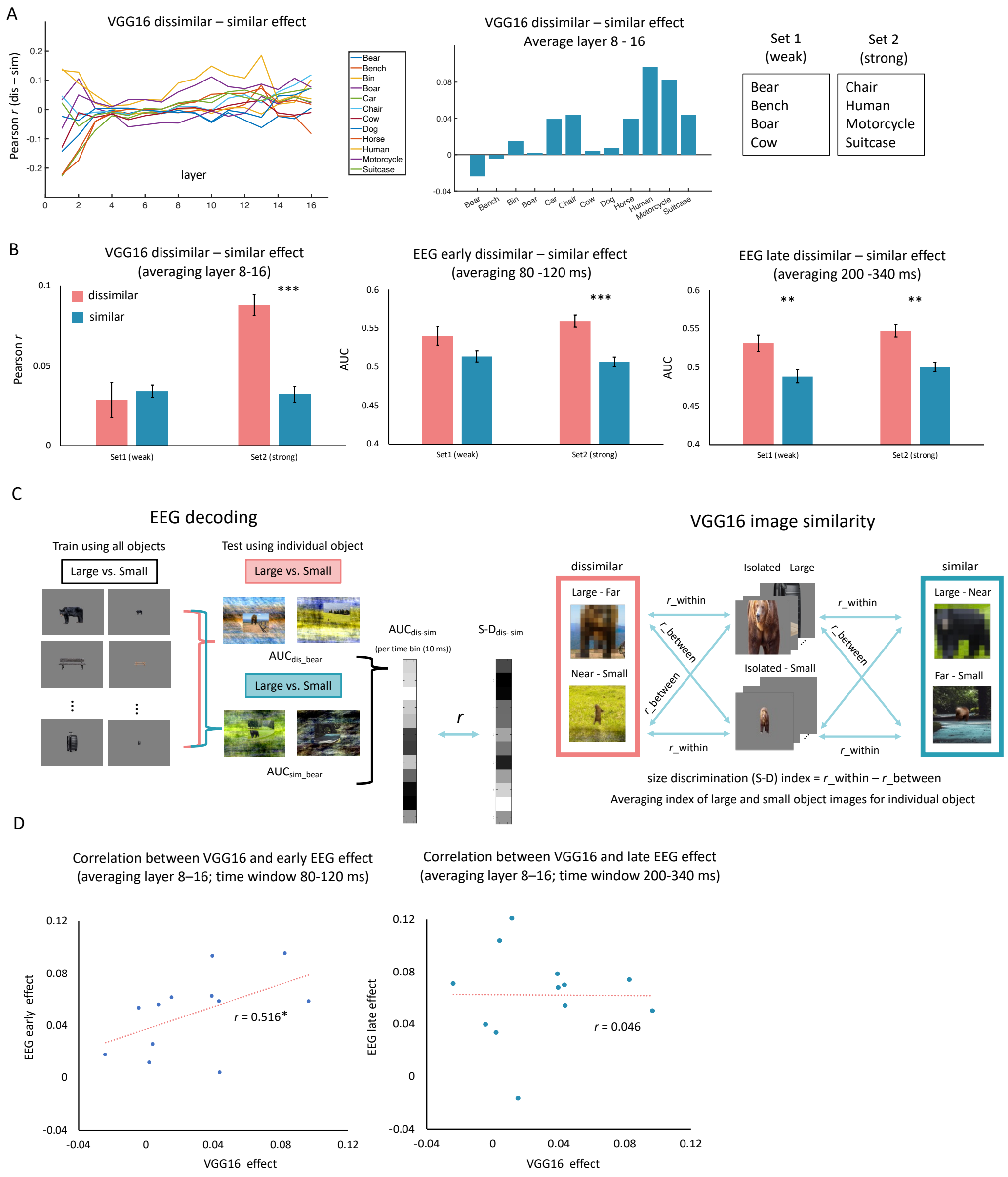


**Supplementary Figure S2.** Relationship between VGG16 (visual angle = 3 degrees) and key EEG results (i.e., the difference in cross-decoding performance between perceptually similar and dissimilar conditions). (A) Size discrimination index (see Figure S1) in VGG16 across layers, depicted separately for each object category (left panel) and averaged across significant layers (8 - 16) (middle panel). The right panel lists the four object categories with the strongest and with the weakest size discrimination index, as was used to reanalyze the EEG data. (B) The results of comparing ‘weak’ (i.e., unconfounded) and ‘strong’ (i.e., confounded) object sets. The left panel shows the VGG16 image analysis results. Only the strong object set shows a significant difference between dissimilar and similar conditions (*** indicates paired *t*-test, *p*-value <0.001, one-tailed), confirming the selection made in Figure S1A. Next, we redid the EEG cross-decoding analyses, using only the “strong” (i.e., confounded) stimulus set, and only the ‘weak’ (i.e., unconfounded) stimulus set. The middle panel shows the results of the *early* size constancy effect in the EEG cross-decoding analysis, averaging the significant time window (80-120 ms) based on the analyses in the complete dataset. Only the strong object set shows a significant difference between dissimilar and similar conditions (*** indicate paired t-test, *p*-value <0.001, one-tailed). Th weak object set does not. The right panel shows the results of the *late* size constancy effect, averaging the time window (200-340 ms) based on the analyses in the complete dataset. In this case, both weak and strong object sets show significant differences (** indicate paired t-test, *p*-value <0.005, one-tailed). (C) The procedure of the time-resolved correlation analysis between VGG16 and the key EEG effect (difference in cross-decoding between perceptually dissimilar and similar conditions). First, we conducted the EEG cross-decoding analysis by training the classifiers on all isolated images, but testing on each individual object separately. Then, we computed the difference between the two perceptual similarity conditions to obtain the critical difference score for each of the 12 object categories, for every time point (10 ms). Next, we correlated these 12 values per timepoint with the size discrimination index from the VGG16 image analysis (see Supplementary Figure 1B caption for details), and tested whether these correlations differed from chance at the group level (i.e., averaged across the 31 participants). These results, confirming that only the early EEG results correlate with the VGG16 image analysis, are displayed in Figure 3E of the main manuscript. (D) Results of group-level correlations, across object categories, between the VGG16 image analysis (average of layers 8-16) and the key EEG results, separately for the early (left panel) and late size-consistency effect (right panel). Taken together, these supplementary analyses demonstrate that the early size-constancy effect (i.e., <120ms) observed in Experiment 2 can be fully accounted for by visual stimulus confounds (as retrieved from the VGG16 image analysis). Crucially, this is not the case for the late size constancy effect (i.e., >200ms).
